# Absence of short-term axon initial segment plasticity in human, mouse, and rat cortical circuits

**DOI:** 10.1101/2025.09.01.672986

**Authors:** Anna Sumera, Laura S Oliveira, Angelika Kwiatkowska, Kirsty Haddow, Rob McGeachan, Lewis W Taylor, Karen Burr, Siddharthan Chandran, Giles Hardingham, Imran Liaquat, Claire Durrant, Paul M Brennan, Peter C Kind, Sam A Booker

**Author notes:** co-corresponding authors, Correspondence to: Sam A Booker, Simons Initiative for the Developing Brain, Centre for Discovery Brain Sciences, Hugh Robson Building, George Square, Edinburgh, EH8 9XD, Scotland, UK,; Peter C Kind, Simons Initiative for the Developing Brain, Centre for Discovery Brain Sciences, Hugh Robson Building, George Square, Edinburgh, EH8 9XD, Scotland, UK. co-first authors (Anna Sumera, Laura S Oliveira).

## Abstract

Maintaining neuronal output with respect to input in the physiological range relies on the ability of neurons to update their responsiveness to inputs dependent on changing activity levels. Termed homeostatic plasticity, the mechanisms that neurons employ to control their responsiveness are varied, and proposed to include structural changes to a key neuronal structure – the axon initial segment (AIS). As the site of action potential initiation, the AIS has been postulated to rapidly change its length in response to increased or decreased cellular and circuit activity. To date, AIS structural plasticity has only been tested in tissue cultures and rodent models. In our current study, we assess the ability of neurons to alter their AIS length over a variety of timescales in *ex vivo* rodent and human brain slices, human neurons derived from induced pluripotent stem cells, and in mice dark-reared during early life; using a combination of electrophysiology and immunohistochemistry. We find no evidence for changes to AIS length following depolarisation for up to 3 hours, despite positive controls confirming modulated activity. However, we do find that neuronal physiological properties are altered by changes in activity – but these are largely independent of action potential initiation associated with the AIS. In summary, we find no evidence supporting a role for AIS structural plasticity in mouse, rat, or human cortical neurons.

## Introduction

How neurons respond to changes to their activity is a critical feature governing how the brain responds to new experiences throughout life. Termed homeostatic plasticity, these mechanisms seek to alter the ability of a neuron to respond to new activity, to maintain a basal level of activity within the physiological range. Such homeostatic plasticity occurs over both short and long time-scales, effecting many aspects of neuronal function, such as intrinsic membrane excitability, neuronal structure, and synaptic strength (Turrigiano, 2011). Altered homeostatic plasticity is believed to contribute to a number of neuropathological states, including epilepsy (Lignani et al., 2020), neurodevelopmental conditions (Booker and Kind, 2021, Kaphzan et al., 2011), and dementia (Styr and Slutsky, 2018). One such form of structural plasticity occurs in the axon initial segment (AIS), with evidence indicating it changes length and position in response to neuronal activity (Evans et al., 2015, Grubb and Burrone, 2010, Grubb et al., 2011, Jamann et al., 2021).

The AIS is a unique structural element of neurons, found to emerge from the soma and occasionally proximal dendrites (Thome et al., 2014, Wahle et al., 2022). This structure ensures neuronal polarity by initiating the outgrowth of the axon, and is the initiation site of action potentials (Leterrier, 2018). The AIS is composed of a unique organisation of cytoskeletal elements (including AnkyrinG and β4-spectrin), is enriched in ion-channels required for rapid ionic conductance (including voltage-activated sodium Na_V_ and potassium Kv channels), and is a site for inhibitory synaptic inputs (Leterrier et al., 2015). Given that the AIS is a site of ion channels enrichment and action potential (AP) generation, its structure has been linked to the efficiency with which a neuron responds to activity (Kole et al., 2008). In primary neuronal cultures from rats and mice, the absolute position and length of the AIS has been shown to change in response to depolarising stimuli (e.g. high extracellular K^+^) or reduced activity (e.g. extracellular tetrodotoxin [TTX]) (Grubb and Burrone, 2010). We recently showed that such plasticity was not observed in acute hippocampal slices from wild-type or the *Fmr1^-/y^* mouse model of Fragile X syndrome, when neurons were depolarised for 3 hours (Booker et al., 2020). Rather, we observed homeostatic compensation to this depolarisation in CA1 pyramidal cells through a reduction in membrane resistance, which was independent of AIS length changes. Contrary to this finding, a recent study suggested that AIS-mediated changes in excitability occur over much shorter time-scales, with plasticity occurring after 1-hour of depolarising stimuli (Jamann et al., 2021). Further evidence for AIS length changes was reported following sustained loss of activity *in vivo* following prolonged dark rearing of mice during the visual critical period (Gutzmann et al., 2014). These findings supported previous work demonstrating sustained homeostatic alteration following visual deprivation through other cellular mechanisms, including altering synaptic strength and number (Desai et al., 2002), composition of synaptic N-methyl D-Aspartate (NMDA) receptors (Philpot et al., 2003, Czepita et al., 1994, Fox et al., 1991, Tongiorgi et al., 2003), and neuronal excitability (Benevento et al., 1992, Brown et al., 2019).

In this study, we hypothesised that the AIS would undergo activity-dependent structural plasticity in both rodent and human neurons on both short and long timescales, leading to altered neuronal excitability. Using a combination of whole-cell electrophysiology and immunohistochemistry of acute brain slices from mice, rats, and humans in which we induced sustained depolarisation we failed to reject the null hypothesis – finding no evidence for AIS plasticity over short time-scales. We further assessed if such plasticity was a product of culturing neurons, which we examined with human induced pluripotent stem cells (iPSCs), in which this plasticity was also absent. Finally, dark-rearing from birth also fails to induce AIS length changes. Overall, we provide compelling evidence across a range of species and experimental modalities that the AIS does not undergo structural plasticity on short timescales; whilst observing other forms of homeostatic plasticity.

## Materials and methods

### Animals

*In vitro* electrophysiological experiments were performed in acute slices from either 14 or 28- 35 day-old wild-type mice (C57/Bl6J^CRL^), 21-day-old wild-type rats (Long Evans Hooded - LEH) or 21-day old rats expressing inducible-GFP under the *Fos* promoter (Cifani et al., 2012). For dark-rearing, mice were housed either from birth, or from 14 postnatal-days (P14) until P28-32. Dark-rearing was performed by housing cages inside an environmentally controlled dark-cabinet, inside a darkroom with a light-shielding curtain. Daily welfare checks performed using infra-red night vision goggles and not red-light, as this has been shown to lead to immediate early gene expression (unpublished findings). Darkness was confirmed by placing a piece of photographic film in the dark cabinet, which was exchanged every 3-days to confirm absence of light. Dark-reared mice were housed in nest groups until collected for experiments. Light reared mice and rats were maintained in 12-hour light/dark cycles. All experiments were performed in accordance with institutional (University of Edinburgh, UK) and UK Home Office guidelines (ASPA: PPL: P2262369). All animals were housed in litter-mate cages of 2-6 animals, and given *ad libitum* access to food and water.

### Human brain slice preparation

All human brain tissue collection was subject to local and regional ethical approval (NHS Lothian: REC number: 15/ES/0094, IRAS number: 165488). Prior to tissue collection, all patients provided written consent for surgical access tissue to be used for research, and for use of anonymised patient information (Edinburgh: NHS Lothian Caldicott Guardian Approval Number: CRD19080). No patient identifying information was stored, and all patient information pseudoanonymised. Human neocortical brain tissue was collected during the resection of more deeply located brain tumours (Supplementary Table 1). We received tissue from 14 individuals, of which 57% had glioblastoma, 36% had gliomas, and 7% had brain metastases. Our data consisted of 36% female and 64% male patients, with a median age of 49 years. Within these patients, 64% of patients had no history of seizures and 36% had recent seizure history and been prescribed pre-operative levetiracetam (1g daily, >3 days).

Human brain slices were prepared as previously described (Wilson et al., 2024). Following surgical opening of the skull and dura, a small piece of cortical tissue was resected *en route* to accessing more deeply sited tumours. This brain tissue was transferred to ice-cold, continually oxygenated HEPES modified artificial cerebrospinal fluid (in mM: 87 NaCl, 2.5 KCl, 10 HEPES, 1.25 NaH_2_PO_4_, 25 Glucose, 90 Sucrose, 1 Na-Pyruvate, 1 Na-Ascorbate, 7 MgCl_2_, and 0.5 CaCl_2_; pH 7.35 with NaOH) and then transported to the laboratory (c.a. 20-40 minutes). Brain tissue was then embedded in 3% agar gel (nominally 30 °C) and then sliced using an oscillating blade vibratome (VT1200S, Leica, Germany). For whole-cell recordings, 300 µm thick slices of cortex were cut and transferred to either a submerged storage chamber containing sucrose-modified ACSF (in mM: 87 NaCl, 2.5 KCl, 25 NaHCO_3_, 1.25 NaH_2_PO_4_, 25 glucose, 75 sucrose, 7 MgCl_2_, 0.5 CaCl_2_, 1 Na-Pyruvate, 1 Na-Ascorbate) warmed to 35 °C and bubbled with carbogen (95% O_2_/5% CO_2_) for 30 min then placed at room temperature.

### Rodent brain slice preparation

Rodent brain slices were prepared as previously described (Oliveira et al., 2021). Mice or rats were anaesthetised with isoflurane, the decapitated and their brains dissected into carbogenated (95%O_2_/5% CO_2_) ice-cold sucrose-ACSF. 400 μm thick coronal brain slices containing either the primary somatosensory (S1) or binocular primary visual (V1B) cortex were cut on a Vibratome (VT1200s, Leica, Germany) in semi-frozen sucrose-ACSF, then stored submerged in sucrose-ACSF warmed to 35°C for 30 min and subsequently at room temperature.

### Short term plasticity

Modulation of neuronal activity leading to potential AIS plasticity was induced according to previous studies (Grubb and Burrone, 2010, Evans et al., 2015, Booker et al., 2020, Jamann et al., 2021). In cell-culture, Neurobasal A media was supplemented with 15 mM KCl or NaCl, from 1 M stocks and cultures returned to the incubator for 3 hours at 37 °C. For acute slice plasticity, following recovery at 35 °C (as above) slices were transferred to a holding chamber containing recording-ACSF, with 15 mM KCl, 15 mM NaCl, 20 μM bicuculline, or 50 μM DL-AP5 and 300 nM tetrodotoxin (TTX) added on top of baseline ionic concentrations. Slices were then incubated for 1 or 3 hours at room temperature (20-22 °C), and carbogenated throughout. Following incubation coverslips or slices were immediately immersion fixed with 4% paraformaldehyde (PFA) in 0.1 M phosphate buffer (PB), pH 7.35 for 20 minutes (coverslips) or 1 hour (slices) at room temperature. For physiology, slices were transferred to the recording chamber circulating with fresh ACSF and further recordings performed.

### Human iPSCs

Neurons were derived from human iPSCs as previously described (Bilican et al., 2014). Neuronal precursor cells (NPCs) were derived from iPSC lines, In order to obtain differentiated neurons, NPCs were then plated in default media (ADMEM/ F12, 1% P/S, 0.5% Glutamax, 0.5% N2, 0.2% B27, 2 mg/mL Heparin) on poly-D-lysine, laminin, fibronectin, and matrigel coated coverslips and fed twice a week, maintained in an incubator at 37 ^°^C, 3% O_2_ & 5% CO_2_. Cultures were supplemented with 10 mM forskolin in weeks 2 and 3 and with 5 ng/mL BDNF and 5 ng/mL GDNF in week 4. For the human neuron/mouse astrocyte co-culture NPCs were plated in DIV4 astrocytes cultures obtained from E17.5 CD1 mouse embryos as previously described (Bell et al., 2015). 5 to 6 week-old hiPSC derived neurons were incubated for 3h at 37 °C in a 5% CO_2_ incubator with 15 mM NaCl or KCl added to the media, after which time cover slips were fixed with 4% PFA for 15 minutes, then transferred to PBS.

### Whole-cell electrophysiology

For electrophysiological recordings, slices were transferred to a submerged recording chamber perfused with pre-warmed carbogenated ACSF (in mM: 125 NaCl, 2.5 KCl, 25 NaHCO_3_, 1.25 NaH_2_PO_4_, 25 glucose, 1 MgCl_2_, 2 CaCl_2_) at a flow rate of 4-6 mL.min^-1^ at 31± 1 °C).Slices were visualised under infrared differential inference contrast microscopy with a digital camera (SciCam Pro, Scientifica, UK) mounted on an upright microscope (SliceScope, Scientifica, UK) and an infinity-corrected 40x water-immersion objective lens (N.A. 0.8, Olympus, Japan). Whole-cell patch-clamp recordings were performed with a Multiclamp 700B (Molecular Devices, CA, USA) amplifier. Recording pipettes were pulled from borosilicate glass capillaries (1.7 mm outer/1mm inner diameter, Harvard Apparatus, UK) on a horizontal electrode puller (P-97 or P-1000, Sutter Instruments, CA, USA). For recordings, pipettes were filled with a K-gluconate based internal solution (in mM 142 K- gluconate, 4 KCl, 0.5 EGTA, 10 HEPES, 2 MgCl_2_, 2 Na_2_ATP, 0.3 Na_2_GTP, 10 Na_2_Phosphocreatine, 2.7 Biocytin, pH=7.4, 290-310 mOsm), resulting in 4-7 MΩ tip resistance. Cells were rejected if: they were more depolarised than -50 mV, initial series resistance >30 MΩ, or series resistance changed by more than 20% over the course of the recording.

All intrinsic membrane properties were measured in I-clamp. Passive membrane properties, including membrane time constant, input resistance, were measured small hyperpolarising steps (10 pA, 500 ms duration), from resting membrane potential. Active properties were determined from a series of depolarising current steps (-100 to +500 pA, 500 ms) from resting membrane potential. All AP properties were determined from the first AP elicited at rheobase. Membrane resonance properties were determined from a chirp stimulus (50 or 100 pA, 20s). sEPSCs were recorded for 1-5 mins in V-clamp. Events were identified using template scaling of a triexponential non-linear regression template based on a subset of events. Only events with amplitude exceeding 3x baseline SD were included.

All recordings were filtered online at 10 kHz with the built-in 4-pole Bessel Filter and digitized at 20 kHz (Digidata1440, Molecular Devices, CA, USA). Traces were recorded in pCLAMP 9 or pClamp10 (Molecular Devices, CA, USA) and stored on a personal computer. Analysis of electrophysiological data was performed offline using the open-source software package Stimfit (Guzman et al., 2014) or using MATLAB.

### Immunohistochemistry and imaging

Immunocytochemistry was performed on free-floating brain slices fixed with 4% paraformaldehyde (PFA) in 0.1 M phosphate buffer (PB) after drug or vehicle incubation and/or following recording; or from sections cut from perfusion fixed brains (for dark-rearing experiments). For perfusion fixation, mice were deeply sedated with isofluorane, then terminally anaesthetised with sodium pentobarbital (50 mg/kg). Mice were then transcardially perfused with room temperature phosphate buffered saline (PBS), then with 20 mL of 4% PFA. Following dissection, brains were post-fixed for 1 hour at room temperature, the transferred to PBS until sectioning. Slices and sections were washed in PBS, and then blocked for 1 hour at room temperature in 10% normal goat serum (NGS), 0.3% TritonX-100, 0.05% NaN_3_ in PBS. Slices and sections were then incubated in primary antibodies raised against Ankyrin G (mouse, 1:500, clone-N106/36 NeuroMab, UNC Davis, CA, USA) and NeuN (rabbit, 1:500, ABN78, Millipore) diluted in PBS containing 5% NGS, 0.3% TritonX-100 and 0.05% NaN_3_, for 24 to 72 hours at 4°C. Slices and sections were thoroughly washed in PBS, then secondary antibodies (anti-mouse and anti-rabbit AlexaFluor488 and AlexaFluor 568, Invitrogen, UK) applied diluted in PBS containing 3% NGS, 0.1% TritonX-100 and 0.05% NaN3, for 3 hrs at room temperature or 24 hrs at 4°C. Samples were rinsed in PBS, then PB and mounted on glass slides with Vectashield hard-set mounting medium (H1400, Vector Labs, UK). For cell-culture blocking time was reduced to 10 minutes; primary antibodies incubated over-night at 4°C and secondary antibody incubation for 1 hour at room temperature. Both primary and secondary antibody solutions were identical to those used in slices, but lacking Triton-X.

Confocal image stacks were collected with an AxiovertLSM 510 (Zeiss, Germany) or SP8 (Leica, German) scanning-confocal microscopes equipped with a 63x (N.A. 1.4, Zeiss, Germany) oil-immersion objective lens. Z-stacks (0.5 µm steps, 1024x1024 pixels) containing ROIs were collected either through the entire 50 µm section (perfusion fixed tissue) or the top 20-30 µm of acute slices. Two stacks of each brain region were collected per experimental condition for each animal. For cell culture experiments, z-stacks (1 µm steps, 1024 x 1024 pixels) were taken from the top to bottom of the monolayer of cells and 2 images per coverslip were collected with a 40x (N.A. 1.3, Zeiss, Germany) oil-immersion objective lens.

### Western-blot analysis

For western blot detection of AnkyrinG, approximately 5 μg of protein from homogenates of V1 were loaded onto a precast gradient gel (3%–8%) and separated by gel electrophoresis using a Tris acetate-based buffer system to ensure maximum resolution of proteins in the molecular weight range of interest. Western blotting onto a PVDF membrane was then performed using the Xcell Surelock system (Invitrogen) according to the manufacturer’s instructions.

Following the protein transfer, the PVDF membrane was blocked for 1 hour at room temperature with 5% (w/v) non-fat dried milk in TBS with 0.1% Tween 20. The membrane was incubated at 4°C overnight with primary antibody raised against Ankyrin G (mouse, 1:500, sc-12719, Santa Cruz Biotechnology) diluted in blocking solution. For visualization of western blots, HRP-based secondary antibodies were used followed by chemiluminescent detection on Kodak X-Omat film. Western blots were digitally scanned and densitometric analysis was performed using ImageJ. All analysis involved normalizing to total protein as a loading control.

### Image analysis

All image analysis was performed with the FIJI package of ImageJ. Based on AnkyrinG or β1-NaV-GFP labelling, AIS were manually traced from their distal tip to the axon hillock, based on AnkyrinG labelling through the 3D image stack using the segmented line tool in FIJI. For fixed tissue, up to 50 AIS were measured for each image, giving a total of up to 100 AIS for each animal or patient. For coverslips, 10 to 15 AIS were measured per coverslip. Independent confirmation of the methodology for AIS measurement was performed by three experimenters, all blind to treatment group, and demonstrated a high degree of consistency. For cFos-GFP fluorescence intensity quantification, ROIs were manually drawn around GFP positive somata, then average fluorescence intensity within each ROI measured. A minimum of 50 somata were counted per condition per rat.

### Statistical analysis

All analysis was performed blind to treatment (where applicable). All data is shown as mean ± SEM. Where applicable, data were analysed with either a linear mixed-effects model (LMM), or its generalised form (GLMM), whereby the variability due to random effects (biological replicate) was taken into account, allowing for direct measurement of treatment effects (Li et al., 2023). Mixed-effects models were fitted using the Lme4 R package (Bates et al., 2014), where the tested variable is a fixed-effect parameter (i.e. species, cortical layeror treatment) and random effects (animal or patient ID) are modelled vectors. All data were tested for normality, with *p*-values reported as the output of Type-3 ANOVA tests. Where reported, statistical significance was assumed if *p* < 0.05.

## Results

The ability of the AIS to undergo structural plasticity on short time-scales has not been consistently observed (Booker 2020, Evans et al., Jamann et al., 2021). To address our main hypothesis that the AIS undergoes structural plasticity following sustained neuronal activation, we performed detailed anatomical and physiological characterisation of the AIS in 3 mammalian species: mice, rats, and humans. A major confound of such analyses is inter-individual variability in response to environmental experience, as such we have sampled from an appropriate number of independent biological replicates based on previous reports. All data is shown as animal averages to prevent over-estimation of effect sizes, but statistically analysed using a multi-compartmental linear-mixed effects model to take into account the high degree of cell-to-cell variability (Booker et al., 2020, Booker et al., 2019, Li et al., 2023, Oliveira et al., 2021).

### The axon initial segment does not display short-term plasticity in mouse, rat and human brain slices

We first tested whether the AIS undergoes short-term synaptic plasticity in acute brain slices in response to changing activity levels in inbred wild-type mice, outbred wild-type rats, and normative human brain tissue. For these experiments, following slicing, sections of neocortex were treated with either 15 mM KCl or 15 mM NaCl osmotic control for 1 or 3 hours, then fixed, and immunohistochemically labelled for AIS (AnkyrinG) and neuronal somata (NeuN). As control, a slice was fixed immediately post-slicing at ∼0 °C and the same labelling performed. We measured AIS associated with neuronal somata in L2/3 of cortex, irrespective of species. For mice and rats, all cortical measurements were taken from somatosensory cortex, as this has been suggested previously to undergo rapid AIS plasticity (Jamann et al., 2021).

In control slices, we observed that the AIS of L2/3 neurons had an average length of 27.5 ± 0.4 µm (in 5 mice), 26.7 ± 1.4 μm (in 7 rats), and 32.0 ± 6.4 μm (in 7 humans) when measured in brain slices fixed immediately after slicing (**Figure 1**). Indeed, the length of L2/3 PC AIS was similar between species (p=0.097, LMM). We next asked if depolarising stimuli, as have been reported to induce AIS shortening previously (Evans et al., 2015, Jamann et al., 2021), resulted in changes in length over short timescales. For this, we treated mouse brain slices with either 15 mM KCl or osmotic control NaCl for 1 or 3 hours, then fixed the tissue and performed immunolabelling (**Figure 1A**). We found that 15 mM KCl did not induce any change in AIS length in mouse brain slices either following 1- (p=0.19) or 3-hour (p=0.65) treatment (**Figure 1B & 1C**). Accordingly, we found no change in the overall distribution of AIS lengths when compared within each biological replicate (**Supplementary Figure 1**). These data are in agreement with our recent findings, also measured in C57/Bl6J mice (Booker et al., 2020).

**Figure 1:**
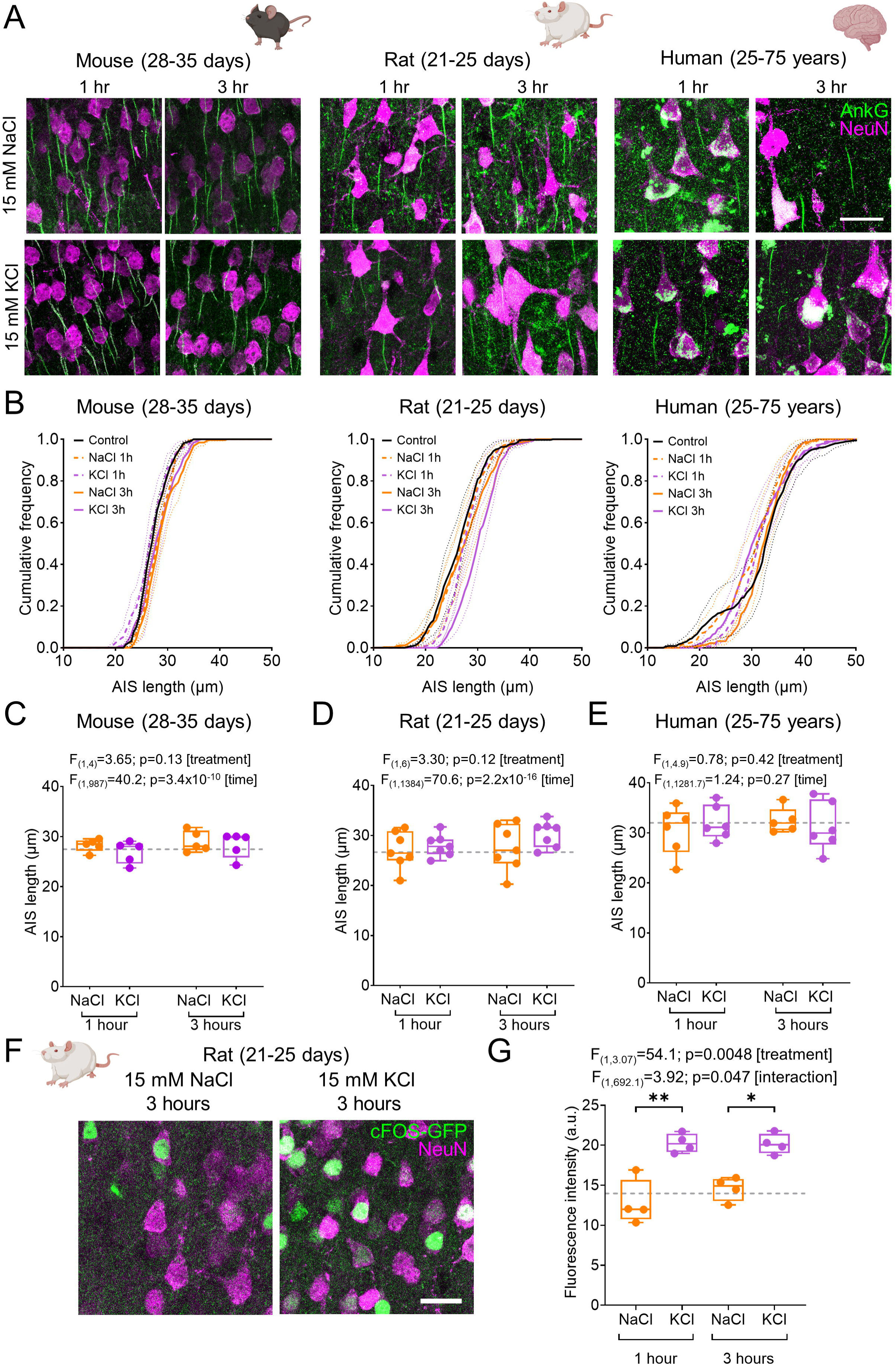
Absence of short-term AIS plasticity following depolarisation in mouse, rat, and human cortex. **A**) immunofluorescence images of AnkyrinG (green) and NeuN (purple) in S1 of P28-35 mice (left) and P21-25 rats (middle), and human neocortex (right). Images shown for 1- or 3-hour treatment with 15 mM NaCl (upper) or 15 mM KCl (lower). Scale bar: 20 μm. **B**) Cumulative frequency plots of AIS length for control (black), and following 1- (dashed lines) or 3-hour treatment (solid lines) with 15 mM NaCl (orange) or 15 mM KCl (purple), shown for S1 of mice (5 mice) and rats (7 rats), and human neocortex (6 patients). **C**) AIS length measured in L2/3 of mouse S1 following 1- and 3-hour treatment with 15 mM NaCl or KCl. Mean AIS length from each mouse (filled circles) and control length (grey dashed line) are shown overlaid. **D**) Average rat S1 L2/3 data, shown in the same format. **E**) Average human L2/3 data, shown in the same format, NaCl 1h: 6 patients; KCl 1h: 6 patients; NaCl 3h: 5 patients; KCl 3h: 6 patients. **F**) Flattened confocal stacks of cFos-induced GFP (green) and NeuN (purple) from L2/3 of S1 treated with either 15 mM NaCl or KCl for 3 hours. **G**) GFP fluorescence (arbitrary units – a.u.) from NeuN positive somata following 15 mM NaCl and 15 mM KCl for 1- or 3-hours. Control GFP fluorescence is shown for reference (grey dashed line). All data is shown as box plots, depicting the median with 25-75% quartile range, and maximum & minimum. Data from individual cells are shown overlaid. Statistics reported from LMM followed by type-3 ANOVA analysis. Some figure elements created with BioRender.

As inbred mice can differ substantially between strains (Lilue et al., 2018) or even between institutes (Wahlsten et al., 2006), we next asked if AIS plasticity is present in outbred rodents, which may take into account a more natural spread of genetic diversity (Tuttle et al., 2018). For this we performed the same experiments measuring AIS lengths in outbred rats. Similar to mice, we found no change in AIS length following either 1 (p=0.81) or 3 hour (p=0.12) administration of 15 mM KCl compared to osmotic controls (**Figure 1B & 1D**). Likewise, we found no change in the distributions of AIS length in each rat (**Supplementary Figure 2**).

Ultimately, we sought to confirm whether AIS shortening occurs in human neurons. To achieve this, we performed identical experiments to rodents, but in acute brain slices prepared from access brain tissue collected during neurosurgical removal of tumours. We collected human data from a total of 14 individuals which had a median age 49 years (Range: 25 – 75 years; see **Supplementary Table 1**). Consistent with our data from rodents, we saw no change in AIS length following either 1 hour or 3-hour administration of 15 mM KCl (Treatment × time: F=2.89, p=0.089) compared to osmotic controls (**Figure 1B & 1E**). Similar to rodents, we saw no consistent change in AIS length distribution when compared within individual (**Supplementary Figure 3**). In human slices, we also measured the AIS lengths of L4 and L5 neurons. Consistent with reports from mice (Gutzmann et al., 2014) we observed layer specific differences in length, such that L2, L3 and L5 had similar AIS lengths (L2 – L3: p=0.89, L2 – L5: p=0.99, L3 – L5: p=0.97; LMM with type 3 ANOVA) and L4 neurons (when present) were consistently shorter (L2 – L4: p=0.049, L3 – L4: p=0.020, L4 – L5: p=0.17; **Supplementary Figure 4**).

One possibility for an absence of AIS plasticity is that our 15 mM KCl treatment failed to induce neuronal depolarisation. To test this, we performed a subset of experiments in a rat line, that expresses GFP under the *Cfos* promoter; such that when cells are activated, they will display increased GFP labelling (Cifani et al., 2012). In brain slices produced from cFos-GFP rats (n=4, **Figure 1F**), we found increased fluorescent labelling of L2/3 cells following 1-hour (p=0.0029) and 3-hour (p=0.012) treatment with 15 mM KCl compared to osmotic controls (**Figure 1G**) and baseline levels (p=0.61 baseline vs. 1 hour, p=0.90 baseline vs. 3 hours). Likewise, treatment of slices with 300 nM tetrodotoxin (TTX) and 50 μM AP-5 to reduce neuronal activity GFP labelling did not differ from baseline (p=0.45, data not shown). These data confirm that 15 mM KCl does increase neuronal activity on short timescales. This indicates that if AIS plasticity were present following 15 mM KCl treatment, our experimental conditions should be suitable to detect it.

Treatment of slices with 15 mM KCl is a profoundly depolarising stimuli (Bianchi et al., 2012), which although reported to induce AIS plasticity, unlikely occurs *in vivo*. To determine if more nuanced modulation of L2/3 PC activity leads to AIS plasticity, we performed identical experiments in brain slices from the mice (n=5, **Figure 2A**) and rats (n=7, **Figure 2B**), by either blocking neuronal activity with TTX (300 nM) and DL-AP-5 (50 μM), or increasing circuit activity by antagonising GABA_A_ receptors with bicuculline (20 μM) to brain slices. In mice, we found that decreasing activity with TTX/AP-5 (p=0.98 vs. baseline) or increasing activity with bicuculline (p=0.93 1-hour, p=0.82 3 hours; vs. baseline) had no effect on AIS length (**Figure 2C**). We observed the same lack of AIS plasticity, when measured in outbred rats, with no effect on AIS length of TTX/AP-5 (p=0.99 vs. baseline) or bicuculline (p=0.93 1-hour, p=0.56 3 hours; vs. baseline; **Figure 2D**).

**Figure 2:**
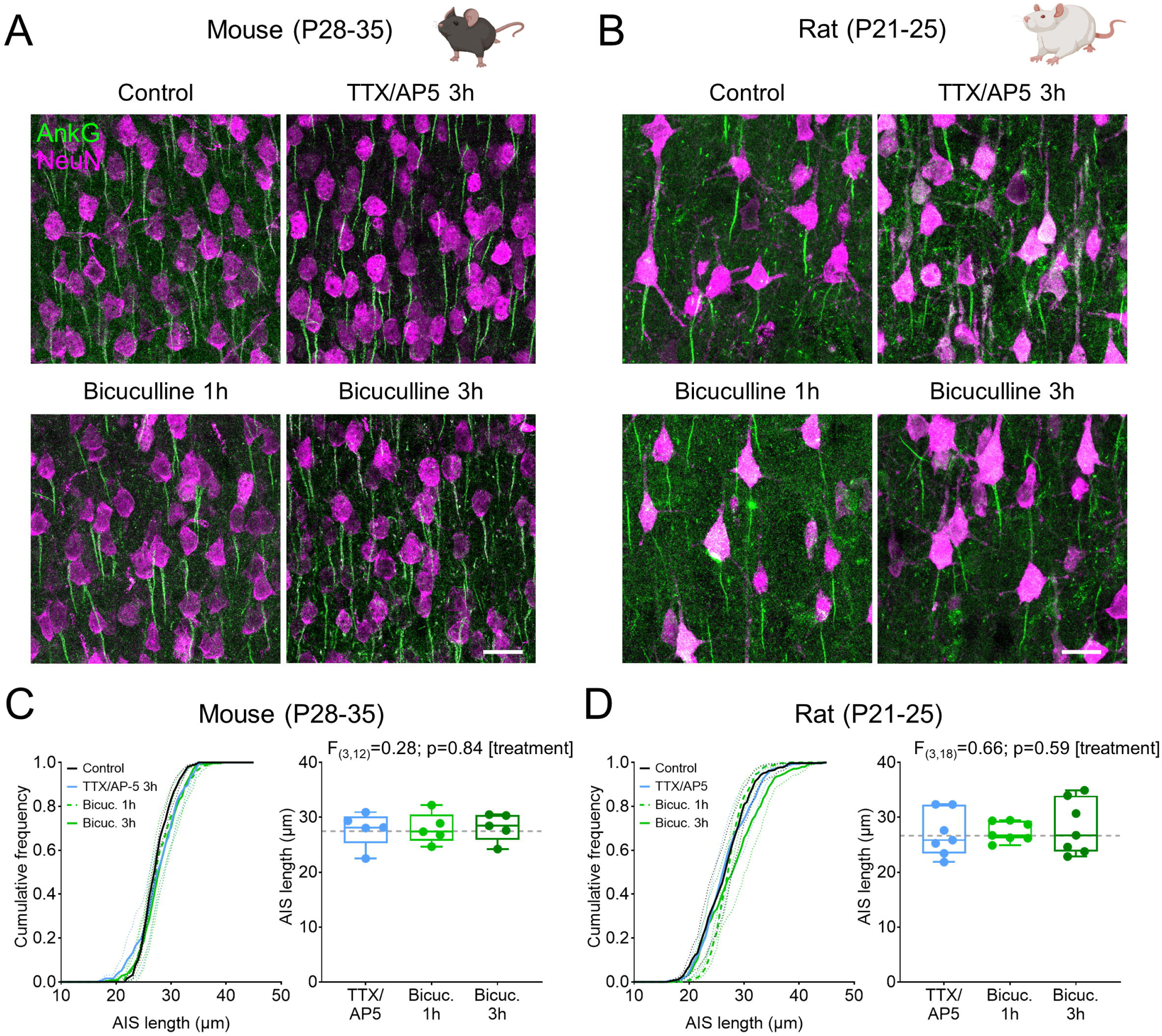
Pharmacological modulation of excitability also does not lead to altered AIS length in mouse and rat somatosensory cortex. **A**) Example flattened confocal stacks of AnkyrinG (green) and NeuN (purple) immunofluorescence labelling in L2/3 of S1 in P28-35 mice, under control conditions, and following 3-hour cessation of neuronal activity with 50 μM DL-AP-5 and 300 nM TTX (AP5/TTX) or following 1- and 3-hour treatment with 10 μM bicuculline methiodide. **B**) Example images from S1 L2/3 of P21-25 rats, according to the same conditions as A. **C**) Left, quantification of AIS lengths as cumulative frequency plots of control (black), TTX/AP5 (blue), 1-hour bicuculline (light green) or 3-hour bicuculline (dark green), n=250 AIS per group from 5 mice. Right, average AIS length for the same conditions compared to control lengths (F=0.28, p=0.84; LMM with type 3 ANOVA). Data from individual mice overlaid. The average control AIS length is shown (grey dashed line). **D**) Quantified AIS lengths in P21-25 rats (n=350 AIS per group from 7 rats) according to the same scheme as C (F=0.66, p=0.59; LMM with type 3 ANOVA). All data is shown as box plots, depicting the median with 25-75% quartile range, and maximum & minimum. Data from individual cells are shown overlaid. Statistics reported from LMM with type-3 ANOVA. Some figure elements created with BioRender.

On balance, our data shows that in acute brain tissue from mice, rats, and humans, that wholesale changes in neuronal excitability over the time-course of 1-3 hours fails to elicit plasticity of AIS length.

### Absence of AIS plasticity in neurons derived from iPSCs, independent of astrocytes

Given the absence of short-term AIS plasticity in brain tissue from adult humans, we next asked if such plasticity is present in human induced pluripotent stem-cell cultures (iPSC), as this has been reported for neurons in primary dissociated cell-culture of mouse brain tissue (Booker et al., 2020, Evans et al., 2015, Sohn et al., 2019). To determine if culture human neurons display similar features, we next performed structural plasticity experiments from iPSC cultures of human neurons alone (**Figure 3A**) or with glial support (**Figure 3D**). In these cultures (n=3-4 independent cultures) we found an average AIS length of cortical neurons derived from iPSCs to be 30.5 ± 1.7 µm in the presence of glia, significantly longer than those in the absence of glia (25.4 ± 1.6 µm, p<0.0001, LMM). In response to 15 mM KCl application for 3 hours, we observed no change in AIS length compared to 15 mM NaCl osmotic controls either cultured alone (p=0.06, **Figure 3C)**, or with glial support (p=0.59, **Figure 3F**). These data confirm that wholesale depolarisation of neurons fails to induce substantial changes in AIS length.

**Figure 3:**
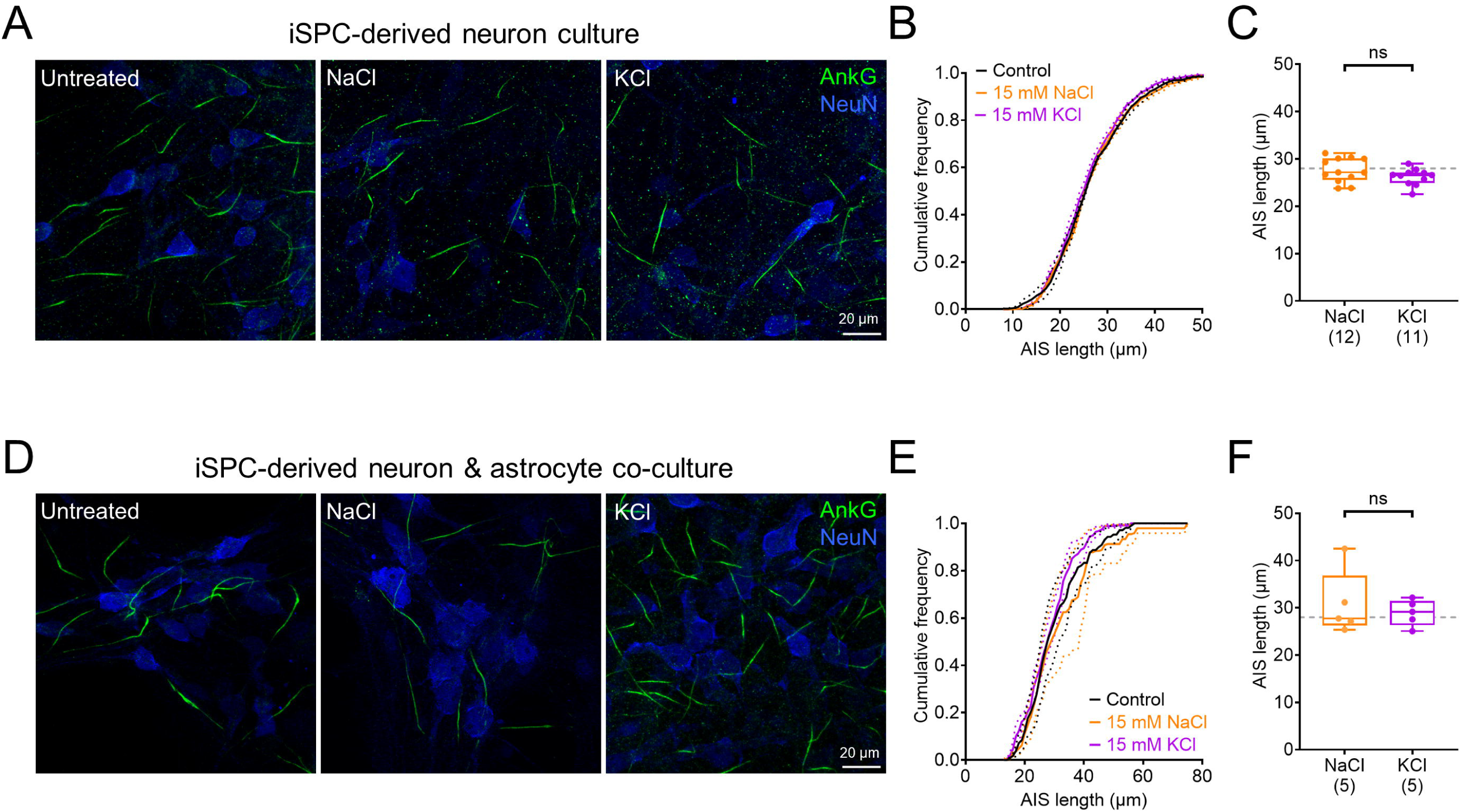
Short-term AIS plasticity is not present in immature human neurons and does not depend on the presence of astrocytes. **A)** representative flattened confocal stacks of human iPSC-derived cortical neurons from cultures treated with either 15 mM NaCl or 15 mM KCl for 3 hours. Neurons are labelled for AnkyrinG (green) and NeuN (red). **B)** cumulative distribution of AIS length from control (black), 15 mM NaCl (orange), and 15 mM KCl (purple) coverslips, data is shown averaged per independent culture. **C)** AIS length measured for each coverslip from 3-4 independent cultures for 15 mM NaCl (orange, n=12 coverslips, 4 cultures) and 15 mM KCl (purple, 11 coverslips, 4 cultures). The average AIS length under control conditions is shown for reference (grey dashed line). **D – F)** the same data but presented for cultures in which iPSC derived cortical neurons were co-cultures with astrocytes. Data are shown for control (black, 6 coverslips from 3 cultures) 15 mM NaCl (orange, 5 coverslips from 3 cultures), and 15 mM KCl (purple, 5 coverslips from 3 cultures). Scale bars: 20 µm. All data is shown as box plots, depicting the median with 25-75% quartile range, and maximum & minimum. Data from individual cells are shown overlaid. All statistics were performed using a LMM with type-3 ANOVA (**C, F**).

### Short-term depolarisation in human L2/3 brain slices reveals AIS length-dependent changes in function

Our anatomical data suggest that short-term depolarisation does not lead to structural plasticity of the AIS. Nevertheless, such changes in neuronal activity have been shown to alter neuronal excitability, such as through modulation of intrinsic membrane properties (Booker et al., 2020, O’Leary et al., 2010) and synaptic function (Ibata et al., 2008, Leslie et al., 2001). To ascertain whether human neurons undergo similar modulation despite no structural changes to the AIS, we performed whole cell patch clamp recordings from L2/3 neurons and measured intrinsic excitability and spontaneous synaptic inputs.

L2/3 neurons in human cortex were recorded from 12 individuals (**Figure 4A** and **Supplementary Table 1**), and typically possessed hyperpolarised resting membrane potentials and accommodating trains of action potentials (**Figure 4B**). Unlike for juvenile rodent hippocampal neurons (Booker et al., 2020), human neurons displayed remarkably few changes in electrophysiological properties in response to sustained depolarisation for either 1- or 3-hours with 15 mM KCl (**Figure 3C**). Contrary to our previous findings in 4- week-old mice (Booker et al., 2020), no change in overall current-frequency slope was observed in human L2/3 neurons (**Figure 4D**, p=0.86, LMM). However, we did observe more depolarised action potential voltage thresholds following 3-hour treatment with 15 mM KCl (**Figure 3E**, p=0.035, LMM), but with no change in rheobase current (**Figure 4F**, p=0.83, LMM) – likely due to concomitant depolarisation of the resting membrane potential (p=0.027, LMM). Previous reports have shown altered synaptic inputs following neuronal activity shifts (Ibata et al., 2008, Jamann et al., 2021), thus we measured spontaneous synaptic inputs to human L2/3 neurons following either NaCl or KCl treatment (**Figure 4G**). Measurement of EPSC events revealed no reduction of spontaneous EPSC frequency in L2/3 PCs, following 3 hours 15 mM KCl treatment (**Figure 4H**, p=0.21, LMM). This was matched by the lack of any observable change in spontaneous EPSC amplitude (**Figure 4I**, p=0.49, LMM). A full summary of all physiological data is shown in **Supplementary Table 2.** These data suggest that paradigms inducing neuronal depolarisation fail to induce AIS plasticity, but with subtle changes in neuronal excitability.

**Figure 4:**
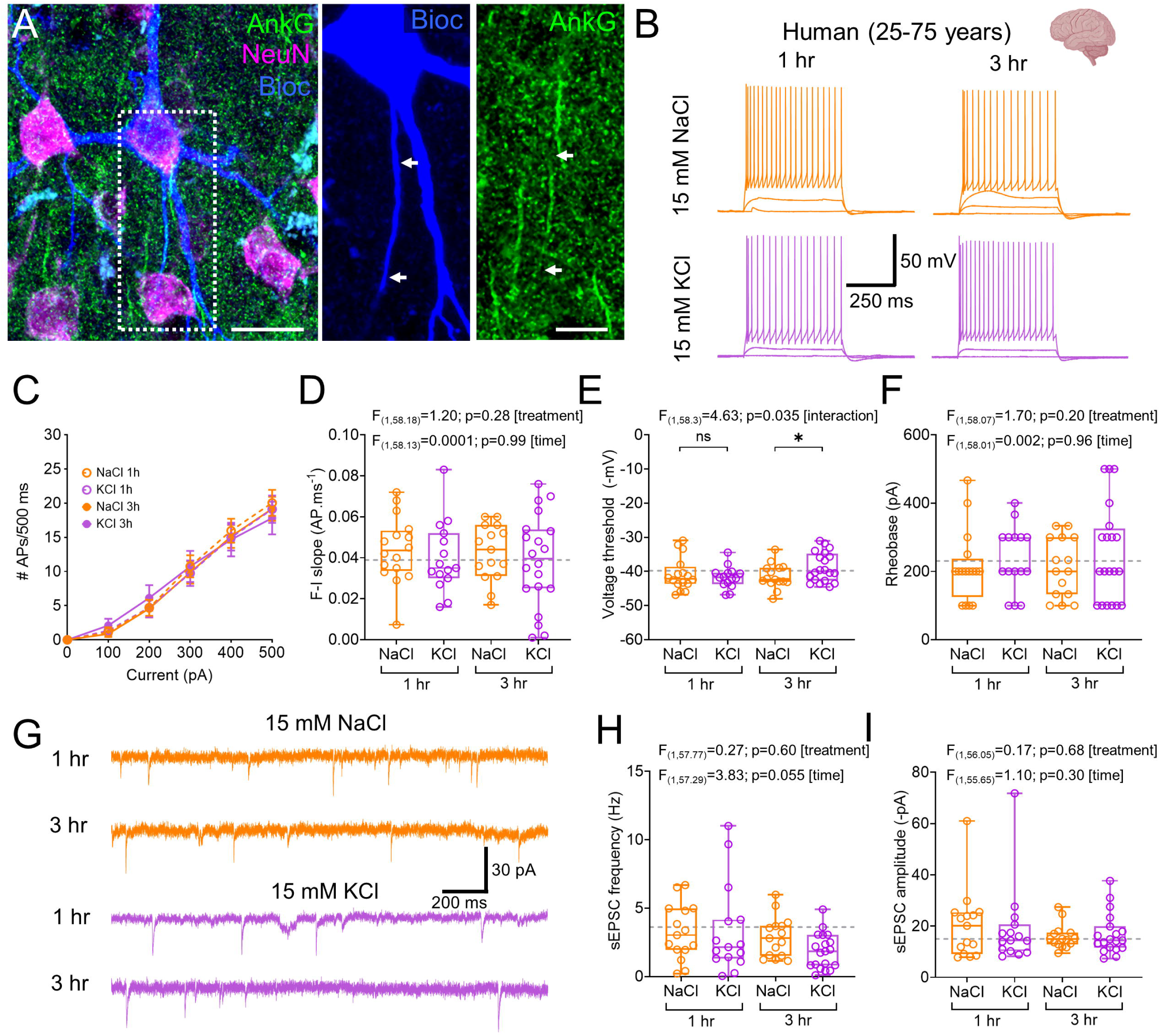
Minor activity-dependent changes in intrinsic neuronal excitability and synaptic function in L2/3 human neurons. **A**) Flattened confocal stack of recorded L2/3 PC in human cortex filled with biocytin (blue), with respect to AnkyrinG (green) and NeuN (purple) immunolabelling. Inset, zoomed in image of the same cells AIS, with the extent of AnkyrinG labelling indicated (arrows). **B**) Voltage response of L2/3 PCs in human cortex to depolarising current steps including maximal firing at 500 pA (100 pA steps, 500 ms duration). Traces are shown for 15 mM NaCl (orange) and 15 mM KCl (purple) following 1- (upper) and 3-hour (lower) treatment. **C**) current-frequency plot for 1-hour (dashed lines) and 3-hour (solid lines) treatment with 15 mM NaCl or KCl. Data shown from: NaCl 1h: n=16 cells from 6 patients; KCl 1h: n=16 cells from 6 patients; NaCl 3h: n= 15 cells from 6 patients; KCl 3h: n=20 cells from 6 patients. **D**) Current-frequency response slope (F-I slope) for recorded cells for all treatment groups. **E**) voltage threshold for the first action potential for all treatment groups. **F**) measured rheobase current for all treatment. **G**) spontaneous EPSC (sEPSC) responses recorded at -70 mV voltage clamp from 15 mM NaCl treatments (upper) and KCl (lower). **H**) measured sEPSC frequency in L2/3 PCs from 15 mM NaCl and KCl for 1- and 3 hours. **I**) measured sEPSC amplitude in L2/3 PCs from 15 mM NaCl and KCl for 1- and 3 hours. All data is shown as box plots, depicting the median with 25-75% quartile range, and maximum & minimum. Data from individual cells are shown overlaid. All statistics were performed using a LMM with type-3 ANOVA. Some figure elements created with BioRender.

Earlier reports in rodents suggest a possible correlation of key cellular excitability features with AIS length, albeit from a small sample size, in a cell specific manner (Jamann et al., 2021). To confirm whether AIS length correlated with neuronal excitability under basal conditions, or following modulation of activity, we measured AIS lengths from a subset of the cells shown in Figure 4, then correlated their length with key intrinsic physiological parameters to allow determination of baseline and excitability-dependent features (**Figure 5**). Under control conditions (**Figure 5A**), we found no correlation between AIS length and rheobase current under basal conditions, indicating that similar current was required to produce action potentials from rest, regardless of axonal structure. However, AIS length did positively correlate with voltage threshold, consistent with the accumulation of sodium channels in this cellular compartment. Somewhat unexpectedly, under basal conditions we found a strong correlation of AIS length with resting membrane potential, but no correlation with input resistance. These baseline data suggest that human neurons from frontal and temporal cortices maintain a constant rheobase as their homeostatic setpoint, which is compensated for by either AIS length or membrane potential, or both. This is in direct opposition to earlier studies that suggest rheobase is the dominant AIS dependent feature (Jamann et al., 2021).

**Figure 5:**
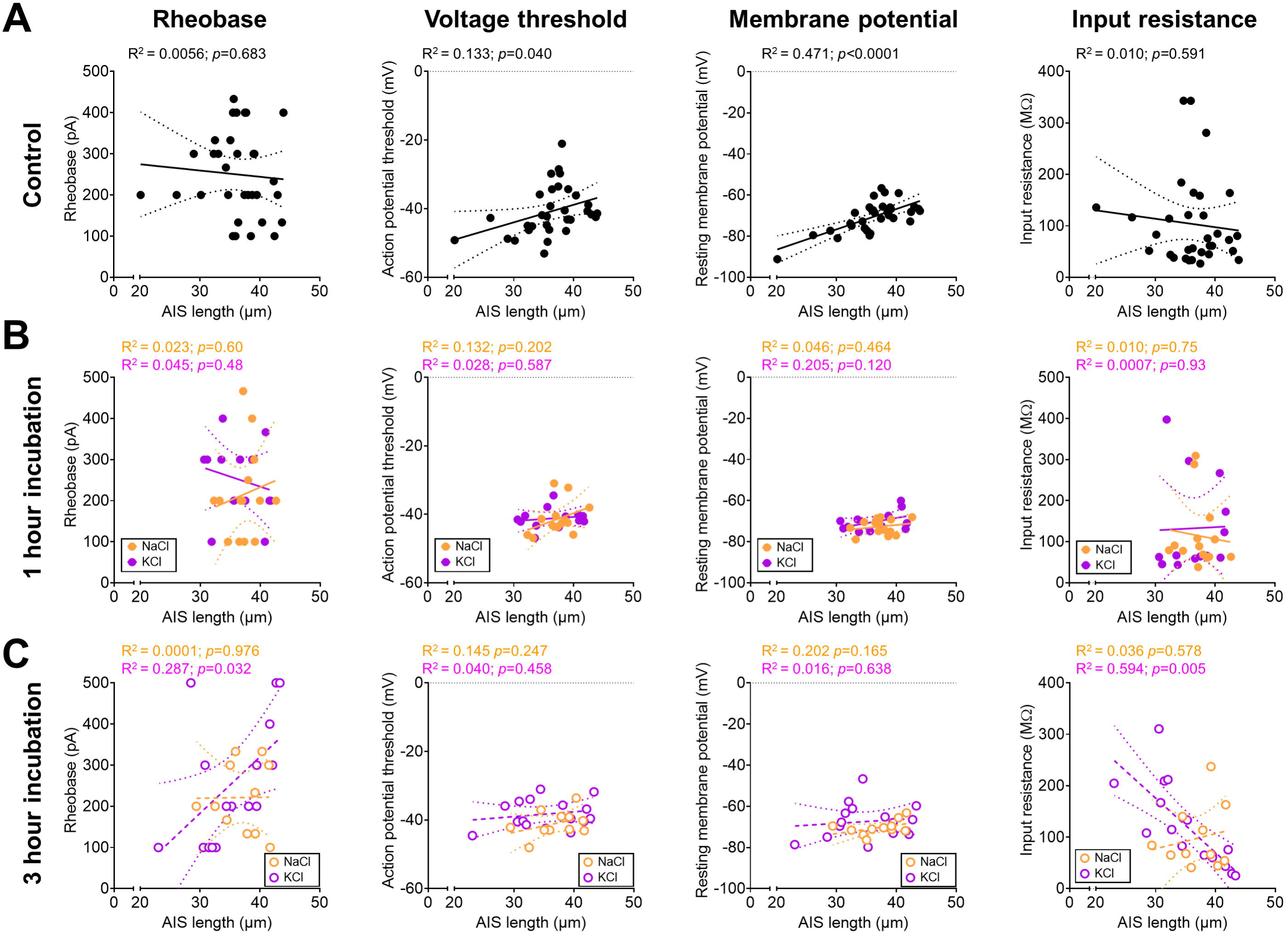
Correlation of AIS length with intrinsic physiology reveals length-dependent modulation of excitability in human L2/3 neurons. **A**) scatter-plots of AIS length (x-axis) versus rheobase, action potential voltage threshold, resting membrane potential, and input resistance for control neurons. **B**) the same data as **A)** but plotted for 15 mM NaCl (orange) and KCl (purple) after 1 hour of treatment. **C**) as for **A)** & **B)** but plotted for 15 mM NaCl (orange) and KCl (purple) after 1 hour of treatment. All data are shown as results from individual cells with linear-regression plotted (dashed line) with the 95% confidence interval (dotted lines). R^2^ and p-values of the regression are shown on the graphs.

To determine whether prolonged depolarisation alters the correlation of AIS length with intrinsic physiology we compared the correlations for 1-hour treatment with 15 mM NaCl or KCl (**Figure 5B**). We found no obvious correlation of any parameter with AIS length under these conditions, contrary to control conditions, although there was a tendency towards AIS length varying with voltage threshold, albeit with a much-reduced sample size. For 3-hour treatment with 15 mM NaCl or KCl (**Figure 5C**) we again found a tendency for AIS length to vary with voltage threshold in the NaCl treated cells, but most interestingly we found that for 15 mM KCl treated slices rheobase showed a very strong correlation with AIS length, which was mirrored by a strong negative correlation of AIS length with input resistance. Such an effect was absent in osmotic controls. We interpret this altered relationship of AIS length with input resistance to reflect that those cells with shortest AIS (thus most hyperpolarised membranes and voltage thresholds) are most responsive to depolarising stimuli, thus regulate their excitability accordingly.

### Long term modulation of sensory inputs fails to regulate AIS length

Given our inability to induce activity-dependent remodelling of the AIS in slice preparations, we next examined an *in vivo* preparation. A number of studies have described the phenomenon that sustained sensory loss is sufficient to lead to homeostatic upregulation in AIS length (Gutzmann et al., 2014, Jamann et al., 2021). Similar to (Gutzmann et al., 2014) and Jamann et al. (2021), we employed dark rearing of mice during the visual critical period (Fox et al., 1991). Specifically, we housed C57/Bl6J^CRL^ mice in total darkness either from birth (P0) or from 14 days postnatally (P14), until P28-35 (**Figure 6A**). Darkness was ensured with a triple light barrier system, with the use of only night-vision goggles for animal welfare checks. At P28 mice were anaesthetised in darkness, and then terminally perfused and immunohistochemistry for AnkyrinG performed, with images containing L2/3 of primary binocular visual cortex (V1B) imaged for AIS lengths (**Figure 6B**). Dark-reared mice were compared to light reared controls perfused at either P14 or P28, which had AIS lengths of 27.6 ± 0.58 μm and 22.9 ± 0.58 μm respectively (p<0.0001, LMM); consistent with the described developmental downregulation of AIS length (Gutzmann et al., 2014). We observed AIS lengths in dark-reared groups (P0-P28: 23.3 ± 0.63 μm; P14-P28: 21.3 ± 0.61 μm) that were comparable to light-reared controls (p= 0.86 [P0-28], p=0.14 [P14-28], LMM, **Figure 6C**). To confirm that AnkyrinG levels were not altered by dark-rearing, a separate cohort of mice were dark reared for 28-days and western-blot analysis for AnkyrinG performed (**Figure 6D**). We found no change in AnkyrinG abundance (as compared to total protein) in dark-reared mice (t_(12)_=0.69, p=0.51, unpaired 2-tailed t-test), consistent with an absence of AIS structural remodelling.

**Figure 6:**
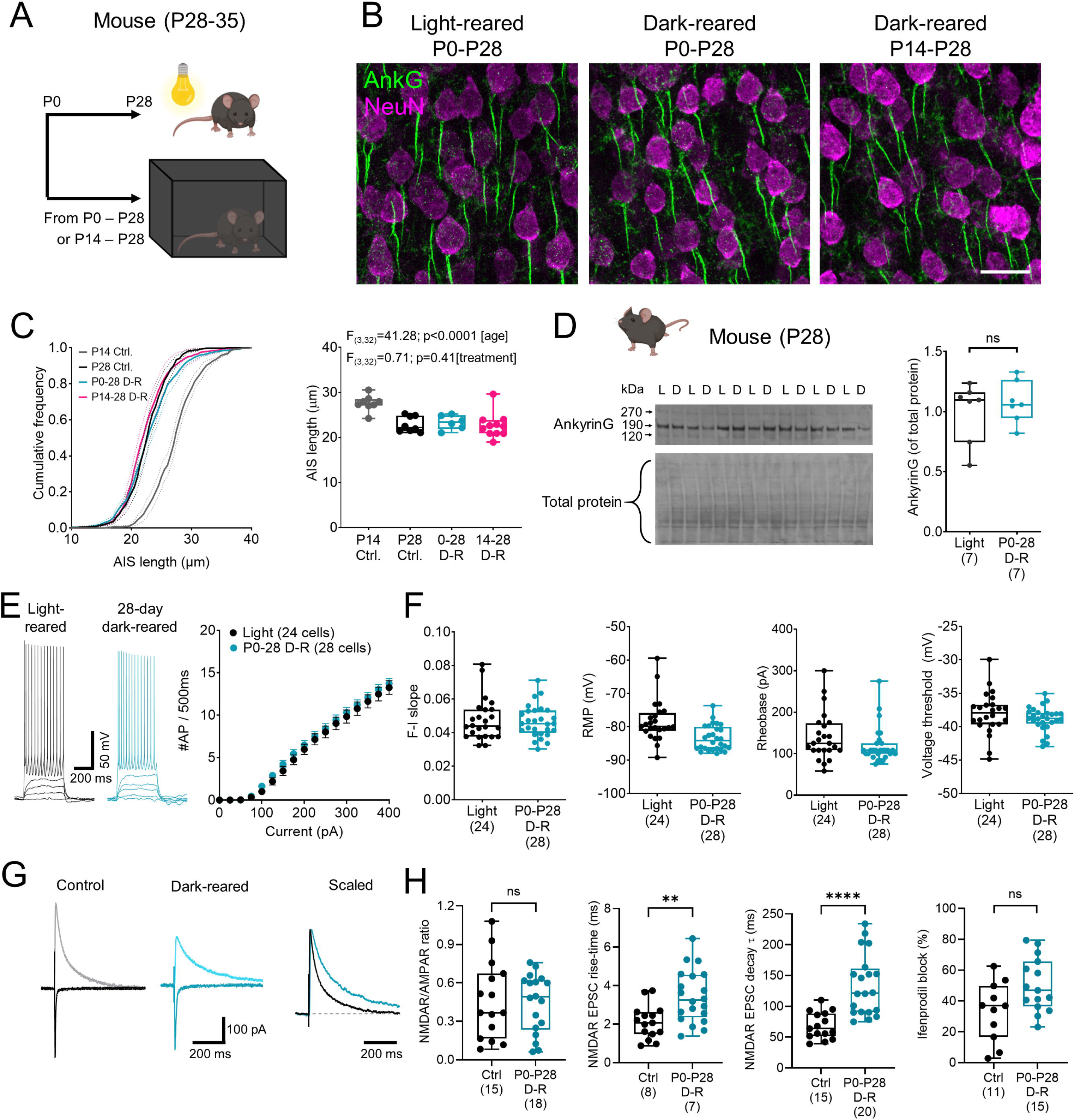
Long-term visual deprivation does not impair AIS length development or neuronal excitability. **A**) schematic of dark-rearing experiment, in which mice are housed in the dark from either birth (P0) or P14. **B**) example flattened confocal stacks of AnkyrinG (green) and NeuN (purple) immunofluorescence labelling in L2/3 of V1B in P28 mice, which have been light-reared (left), dark-reared from P0-P28 (middle) or dark-reared from P14-P28 (right). Scale bar: 20 μm. **C**) Cumulative frequency plots of all AIS lengths measured for each mouse and averaged within group: P14 light-reared (grey), P28 light-reared (black), P0-P28 dark-reared (0-28 D-R; teal) and P14-P28 dark-reared (14-28 D-R; magenta). Right, average AIS lengths for each mouse plotted according to treatment group. **D**) Western-blot analysis of V1 from light-reared and P0-28 D-R mice. Example blots for AnkyrinG (upper) and total protein (lower) are shown from light-reared (L) and dark-reared (D) mice. Right, Quantification of AnkyrinG intensity normalised to total protein. **E**) example voltage response of L2/3 PCs in mouse V1B in response to depolarising current steps including maximal firing at 400 pA (25 pA steps, 500 ms duration) from light-reared controls (black) and mice dark-reared from P0-P28 (teal). **F**) current-frequency plots of L2/3 V1B neurons in mice from light- (24 cells from 7 mice) and dark-reared (28 cells from 8 mice) experiments. **G**) quantification of F-I slope of the same cells. **H**) resting membrane potential of the same cells. I) rheobase of the same cells. J) action potential voltage threshold of the same cells. All data is shown as box plots, depicting the median with 25-75% quartile range, and maximum & minimum. Data from individual cells are shown overlaid. All statistics were performed using a LMM with type-3 ANOVA. Some figure elements created with BioRender.

To confirm whether neuronal function was altered despite a lack AIS length change, we performed whole-cell patch-clamp recordings from L2/3 PCs in V1B of light- and dark-reared (P0 to P28-35) mice. L2/3 PCs in both groups of mice produced reliable action potential discharge in response to depolarising stimuli (Figure 6F), which showed no dark-rearing dependent differences in overall current-frequency response (F_1,50_=0.47, p=0.5, 2-way ANOVA, **Figure 6E**). Accordingly, we observed no difference in key measures of excitability (**Figure 6F**), chiefly current-frequency response slope (p=0.64, LMM), rheobase (p=0.17, LMM) or voltage threshold (p=0.38, LMM). However, we observed a small but significant change in resting membrane potential (RMP) towards a more hyperpolarized RMP in cells from dark reared animals (Ctr: -78.3 ± 1.05 mV, P0-28 DR -83.4 ± 0.96 mV, p=0.04, LMM). A complete summary of all intrinsic parameters is shown in **Supplementary Table 3**. Synaptic NMDAR-mediated currents are known to display delayed maturation following dark-rearing from birth (Czepita et al., 1994, Fox et al., 1991, Tongiorgi et al., 2003). To confirm this effect was present in our sample, we performed electrical stimulation of L1 in V1B while recording from L2/3 PCs in light- and dark-reared mice (**Figure 6G**). We found that while P0- 28 dark-rearing of mice had no effect on overall NMDAR/AMPAR ratio (P=0.78, LMM), it significantly prolonged NMDAR-EPSC rise and decay kinetics (p=0.004 & <0.001 respectively, both LMM), and a tendency towards greater ifenprodil block of pharmacologically-isolated NMDAR EPSCs (p=0.09, LMM). Together, these data support our conclusion that while dark-rearing does modulate synaptic function in V1, it does not lead to changes in AIS length. These data question the reproducibility of AIS plasticity following sensory deprivation, and degree to which it is biologically relevant.

## Discussion

In the present study we found no evidence for AIS length changes following sustained depolarisation up to 3 hours in both rodent and human cortices. Depolarisation modestly modified the excitability of adult human neurons which correlated with AIS length but not action potential threshold. Our data question the reproducibility of AIS structural plasticity and the extent it contributes to cortical function.

### Absence of short term AIS plasticity in cortical circuits

In agreement with our previous findings (Booker et al., 2020), we provide multiple lines of evidence in human neurons that AIS plasticity may not be as consistent a mechanism of homeostatic compensation as previously described (Evans et al., 2015, Grubb et al., 2011, Gutzmann et al., 2014, Jamann et al., 2021). Indeed, we find that although AIS length does correlate with many features of neuronal excitability (especially following depolarising stimulation) these features do not correlate with AP initiation. Nevertheless, we find a strong correlation of AIS length with AP voltage threshold – consistent with previous reports (Kole et al). However, compared with recent reports (Jamann et al., 2021), we find no correlation of AIS length in human cortex with the rheobase current required for AP discharge. This may be due to the strong correlation between resting membrane potential and AIS length we observed, which likely offsets the AP threshold correlation – leading to stabilisation of rheobase current – and reflecting true homeostatic control of neuronal discharge. This may indeed reflect species-dependent differences in AIS function (Libé-Philippot et al., 2023); however, immunolabelling of our previously recorded neurons did not reveal any notably labelling for LRRC37B (a human specific marker of AIS subtypes) – precluding further sub- classification of our data. The principal changes observed in our data following sustained depolarisation were reduced input resistance, consistent with altered leak K^+^ conductance – as previously described in rodent primary cultures and brain slices (O’Leary et al., 2010, Booker et al., 2020). Another important consideration for differences between our current study, and recent efforts to measure AIS plasticity, is the use of appropriate controls. Indeed, Jamann et al. (2021) did not account for time-matched controls when examining the mechanisms of short-term AIS plasticity, which may mask an absence of biological effect. Our presented data are compared to osmotic and time-matched controls, with intra-animal effects variability accounted for in our statistical model, thus reducing the risk of type-1 statistical errors and incorrect rejection of our null hypothesis.

### Absence of activity dependent shortening and ramifications for circuit fidelity

We provide further evidence for an absence of AIS plasticity through the use of dark-rearing. Our data fail to reject the null-hypothesis that loss of visual input does not alter AIS length, following either 2- or 4-week visual deprivation. Despite this, we do observe the described developmental shortening of AIS in V1, over the 2^nd^ to 4^th^ weeks of life, consistent with previous observations (Gutzmann et al., 2014). These data confirm that the AIS is capable of modifying its length as activity patterns in the cortex emerge and mature. It is thus puzzling that loss of visual inputs doesn’t lead to AIS length changes when measured after dark rearing. There are several possible explanations for this apparent discrepancy is activity- dependent control of AIS length: 1) neurons compensate their excitability through other mechanisms to ensure a maintained rheobase, 2) that V1 neurons are not singly responsive to visual inputs and thus loss of light alone is not sufficient to prevent AIS shortening, or 3) that AIS length changes are directly regulated by genetic cues during development. Our current data suggests the former, as we observed no major change in the intrinsic electrophysiology of dark-reared mice compared to control, but as we only performed limited measurement other modifications (e.g. synaptic summation) cannot be excluded (Pandey et al., 2022). However, it is known that V1 in rodents encodes many other salient behavioural features other than just vision (Murray et al., 2016). Indeed, in rodents L2/3 PCs and interneurons produce robust responses to movement (Pakan et al., 2016), somatosensation (Beauchamp et al., 2009) and sound (Williams et al., 2023), given the absence of visual cues to both hemispheres of the cortex it is unclear how much these other sensory modalities modify V1 neuron function and compensate neuronal activity. Whether such compensations occur in experimental models of unilateral sensory deprivation, such as monocular deprivation (reviewed in Gainey and Feldman (2017)) or whisker trimming (Jamann et al., 2021) remains uncertain, but indicates a more complex situation that likely involves re-setting of the homeostatic set-point to maintain activity within the physiological range (Pandey et al., 2022, Bear, 2003).

Activity is not uniform across the population of cortical neurons, thus AIS length changes might not be predicted to be equivalently uniform. This is further supported by the inherent variability of AIS length, which as we (Booker et al. (2020) and current data) and others (Jamann et al., 2021, Kole et al., 2008), but which correlates with biophysical properties of neuronal excitability. Such data strongly support the notion that there is not a “one-size-fits-all” AIS length, and that sampling a small number of neurons from a minimum of biological replicates likely masks biological diversity. Our data reveals a relationship between AIS length and membrane potential in human L2/3 neurons, a duality which may serve to ensure neuronal discharge within the physiological range (Wefelmeyer et al., 2016, Zhao et al., 2024). Such a relationship will ensure that even if neurons receive persistently high activity, they are entrained to firing rates within the physiological range. Our current data shows that following sustained depolarisation, membrane resistance appears to a more profound modifier of neuronal excitability (Shah, 2018), leading to reduced rheobase in neurons that typically fire have hyperpolarised voltage thresholds.

These data have potentially important ramifications for conditions where modified resistance of neurons is a major contributor to neuronal excitability, including neurodevelopmental disorders such as Fragile X Syndrome (Booker et al., 2019, Domanski et al., 2019, Gibson et al., 2008) and SynGAP haploinsufficiency (Michaelson et al., 2018), but also for those in which AIS length changes have been implicated – such as mouse models of Fragile X Syndrome (Booker et al., 2020) and Angelman’s Syndrome (Kaphzan et al., 2011, Liu et al., 2025). If altered membrane resistance and excitability is a crucial factor in rapid cellular responses to elevated excitability, correcting such an effect may short-circuit a neurons ability to regulate activity, which may indeed have unintended consequences for long term brain function.

### Limitations

Our study demonstrates that short term AIS structural plasticity is absent in L2/3 of the neocortex in mice, rats, and humans. This is in opposition with a number of studies finding activity-dependent AIS changes in in acute brain slices of mice (Jamann et al., 2021), organotypic brain slices (Thome et al., 2025), primary cell culture (Evans et al., 2015) and *in vivo* (Gutzmann et al., 2014, Schlüter et al., 2017), but in agreement with our recent work (Booker et al., 2020). One possible key reason for such a discrepancy could be the strain of mouse used. Indeed, multiple lines of evidence now show that inbred mouse lines regularly acquire deleterious mutations leading to hyperexcitability (Rubinstein et al., 2015) and altered GABA receptor function (Mulligan et al., 2019), which can impair circuit function *(Çalışkan et al., 2023)*. Our data seeks to overcome these confounds by performing similar analysis in parallel between multiple species (both inbred and outbred), as well as living human brain tissue. Our data indicates that short-term AIS plasticity following depolarisation is likely not a consistent mechanism of cellular homeostasis across the cortical neuron population.

Our data cannot preclude the possibility of other activity dependent changes in function. For example, recent studies have suggested that GABAergic inhibition selectively targeting the AIS undergoes developmental plasticity and can shape the excitation state of neurons in the absence of AIS structural changes (Pan-Vazquez et al., 2020, Wefelmeyer et al., 2015, Zhao et al., 2024). Our human data cannot address this possible form of plasticity, due to the older age of the majority of the patients we collected tissue from. Future studies should examine the GABAergic synaptic inputs to the AIS in human neurons, as such a feature may be a more reliable mechanism (alongside neuronal resistance changes) to regulate neuron function within active circuits.

In conclusion, we provide a well powered and controlled dataset that confirms the absence of AIS length changes in response to sustained changes in neuronal activity on short time scales. Our data reveal a novel relationship between AIS length and other homeostatic plasticity mechanisms, suggesting a more complex relationship that previously understood. These data have direct implications for how we interpret changes in neuronal function in circumstances of elevated neuronal activity.

## Acknowledgments

Most of all we would like to thank the patients who gave consent for this study, the NHS staff, EMERGE nursing team, and Alsadeg Bilal, for providing time and energy to facilitate tissue collection. We also thank Kind Lab members for helpful discussions. This project was funded through generous support from the Simons Initiative for the Developing Brain (SFARI grant: 529085), The Patrick Wild Centre (AS, SAB & PK), Medical Research Scotland (AK & SAB) the NeuroResearchers Fund (SAB, Edinburgh Neuroscience), RS McDonald Seedcorn Fund (SAB), Dyson Foundation (CD), Race Against Dementia (CD - ARUK-RADF-2019a-00), Alzheimer’s Society (CD-AS-PG-21-006)

## Author contributions

**Anna Sumera:** Investigation, Methodology, Software, Formal analysis; Visualization, Writing - Original Draft, Writing - Review & Editing**; Laura Oliveira**: Investigation, Methodology, Formal analysis; Writing - Original Draft, Writing - Review & Editing, Software; **Angelika Kwiatkowska**: Investigation, Methodology, Software, Formal analysis; **Kirsty Haddow**: Investigation, Methodology, Software, Formal analysis; **Rob McGeachan:** Methodology; **Lewis W Taylor**: Methodology; **Karen Bell**: Resources, Methodology, **Siddharthan Chandran**: Resources, Supervision**; Giles Hardingham:** Resources, Supervision; **Imran Liaquat**: Resources, Methodology; **Claire Durrant**: Resources, Project administration, Writing - Review & Editing; **Paul M Brennan**: Resources, Methodology, Project administration, Writing - Review & Editing; **Peter C Kind**: Conceptualization, Supervision, Funding acquisition, Writing - Review & Editing; **Sam A Booker**: Conceptualization, Investigation, Methodology, Validation, Formal analysis, Visualization, Writing - Original Draft, Writing - Review & Editing, Project administration, Funding acquisition.

## Data availability

All data presented in the figures and text are available in Supplementary Materials. Any further data will be made available upon request.

