## Supplementary materials for "Absence of short-term axon initial segment plasticity in human, mouse, and rat cortical circuits"

*Sumera, Oliveira, et al.*

Supplementary materials:

| **Age range** | **Sex** | **Diagnosis** | **Hemi.** | **Region** | **Seizures?** |
| --- | --- | --- | --- | --- | --- |
| 25-30 | F | Glioma | Right | Frontal | No |
| 30-35 | M | Glioma | Left | Frontal | No |
| 35-40 | F | Glioma | Left | Frontal | Yes |
| 35-40 | F | Glioma | Right | Frontal | No |
| 40-45 | M | Glioblastoma | Right | Frontal | Yes |
| 45-50 | F | Glioma | Right | Temporal | Yes |
| 45-50 | F | Glioblastoma | Right | Frontal | Yes |
| 50-55 | M | Glioblastoma | Right | Frontal | No |
| 50-55 | M | Glioblastoma | Right | Frontal | No |
| 55-60 | M | Metastases | Right | Frontal | No |
| 65-70 | M | Glioblastoma | Left | Temporal | No |
| 70-75 | M | Glioblastoma | Right | Frontal | No |
| 70-75 | M | Glioblastoma | Left | Frontal | No |
| 70-75 | M | Glioblastoma | Right | Frontal | Yes |

Supplementary Table 1: Summary of patients included in the current study, data are shown for age rounded to the nearest 5 years to ensure patient anonymity, biological sex, diagnosis, hemisphere, brain region, and seizure history.

| Electrophysiological Property | Control  (54 cells from 13 patients) | 15 mM NaCl  1 hr treatment  (16 cells from 6 patients) | 15 mM KCl  1 hr treatment  (16 cells from 6 patients) | 15 mM NaCl  3 hr treatment  (15 cells from 6 patients) | 15 mM KCl  3 hr treatment  (20 cells from 6 patients) | LMM (Treatment) | | LMM (Time) | | LMM (interaction) | |
| --- | --- | --- | --- | --- | --- | --- | --- | --- | --- | --- | --- |
|  |  |  |  |  |  | F | p | F | p | F | p |
| Membrane potential (mV) | -69.70 ± 1.03 | -72.04 ± 0.94 | -69.56 ± 1.27 | -69.92 ± 1.34 | -67.77 ± 1.83 | 5.17 | **0.027** | 1.54 | 0.22 | 0.0047 | 0.95 |
| Input resistance (MΩ) | 111.24 ± 11.94 | 157.87 ± 45.70 | 149.97 ± 30.04 | 124.91 ± 28.05 | 112.31 ± 16.54 | 0.35 | 0.56 | 1.33 | 0.25 | 0.065 | 0.80 |
| Membrane time-constant (ms) | 16.31 ± 0.78 | 18.57 ± 2.32 | 15.95 ± 1.46 | 16.35 ± 1.70 | 14.37 ± 1.27 | 2.22 | 0.14 | 1.88 | 0.18 | 0.38 | 0.54 |
| Capacitance (pF) | 203.26 ± 14.45 | 162.09 ± 16.62 | 145.89 ± 15.92 | 172.00 ± 21.15 | 180.99 ± 29.85 | 0.054 | 0.82 | 0.48 | 0.49 | 0.027 | 0.87 |
| Rheobase (pA) | 229.54 ± 13.75 | 213.54 ± 25. 87 | 229.17 ± 24.51 | 211.10 ± 22.46 | 251.67 ± 31.39 | 1.70 | 0.20 | 0.0023 | 0.96 | 0.045 | 0.83 |
| Voltage threshold (mV) | -39.15 ± 0.98 | -40.83 ± 1.21 | -41.85 ± 0.77 | -41.41 ± 0.92 | -38.76 ± 0.96 | 2.09 | 0.15 | 0.88 | 0.35 | 4.64 | **0.035** |
| Peak firing (#APs) | 18.30 ± 1.21 | 19.51 ± 2.06 | 19.10 ± 2.00 | 19.18 ± 1.76 | 17.82 ± 2.39 | 0.58 | 0.45 | 0.062 | 0.80 | 0.030 | 0.86 |
| AP amplitude (mV) | 119.24 ± 1.61 | 120.99 ± 1.43 | 117.59 ± 3.06 | 118.09 ± 2.39 | 117.99 ± 2.39 | 2.10 | 0.15 | 1.03 | 0.31 | 0.20 | 0.66 |
| AP 20-80% rise-time (ms) | 0.16 ± 0.0035 | 0.17 ± 0.0061 | 0.18 ± 0.010 | 0.17 ± 0.0079 | 0.16 ± 0.0036 | 0.45 | 0.50 | 1.55 | 0.22 | 0.92 | 0.34 |
| AP half-height width (ms) | 0.71 ± 0.015 | 0.68 ± 0.028 | 0.71 ± 0.059 | 0.65 ± 0.028 | 0.66 ± 0.022 | 0.25 | 0.62 | 2.20 | 0.14 | 0.034 | 0.85 |
| AP maximum rise (mV.ms^-1^) | 431.94 ± 13.13 | 427.40 ± 18.16 | 411.71 ± 23.60 | 421.79 ± 25.33 | 433.41 ± 13.83 | 0.29 | 0.59 | 0.12 | 0.73 | 0.091 | 0.76 |
| AP maximum decay (mV.ms^-1^) | 116.33 ± 5.54 | 116.67 ± 3.88 | 113.03 ± 4.51 | 124.91 ± 4.30 | 123.11 ± 6.86 | 0.26 | 0.61 | 3.10 | 0.083 | 0.022 | 0.88 |
| IF Slope (AP. pA^-1^) | 0.039 ± 0.0025 | 0.044 ± 0.0041 | 0.41 ± 0.0042 | 0.043 ± 0.0035 | 0.038 ± 0.0050 | 1.20 | 0.28 | 0.0001 | 0.99 | 0.030 | 0.86 |
| Sag (mV) | -3.61 ± 0.51 | -4.22 ± 0.56 | -3.85 ± 0.41 | -3.76 ± 0.34 | -3.38 ± 0.53 | 0.59 | 0.45 | 0.59 | 0.45 | 0.0063 | 0.94 |
| Sag (% of max) | 14.18 ± 1.28 | 12.93 ± 1.42 | 13.30 ± 1.59 | 13.88 ± 1.66 | 12.24 ± 1.55 | 0.19 | 0.67 | 0.0098 | 0.92 | 1.21 | 0.28 |
| Resonant frequency (Hz) | 1.80 ± 0.18 | 1.23 ± 0.32 | 0.86 ± 0.17 | 1.22 ± 0.31 | 1.15 ± 0.30 | 0.28 | 0.60 | 0.16 | 0.69 | 0.17 | 0.68 |
| Peak impedance (MΩ) | 117.90 ± 10.80 | 95.26 ± 14.77 | 96.83 ± 15.96 | 90.94 ± 15.22 | 100.02 ± 14.31 | 0.35 | 0.56 | 0.10 | 0.75 | 0.048 | 0.83 |
| Q-factor | 1.18 ± 0.030 | 1.17 ± 0.074 | 1.33 ± 0.071 | 1.26 ± 0.072 | 1.29 ± 0.050 | 2.86 | 0.097 | 0.21 | 0.65 | 1.06 | 0.31 |
| sEPSC amplitude (pA) | -15.26 ± 1.15 | -20.35 ± 3.50 | -19.38 ± 3.79 | -15.85 ± 1.23 | -16.89 ± 1.83 | 0.17 | 0.68 | 1.10 | 0.30 | 0.48 | 0.49 |
| sEPSC frequency (Hz) | 3.56 ± 0.39 | 3.24 ± 0.53 | 3.48 ± 0.80 | 2.89 ± 0.38 | 1.86 ± 0.30 | 0.27 | 0.60 | 3.84 | 0.055 | 1.64 | 0.21 |
| mAHP (mV) | -15.47 ± 0.61 | -15.16 ± 0.96 | -16.73 ± 0.98 | -15.73 ± 0.69 | -15.72 ± 0.80 | 1.01 | 0.32 | 0.44 | 0.51 | 1.00 | 0.32 |

Supplementary table 2: Summary of measured electrophysiological properties of recorded L2/3 human neurons. All data is shown as mean ± SEM from all experimental groups. Statistics are shown comparing 15 mM KCl and NaCl based on treatment, time, and interaction from LMM analysis.

| Electrophysiological Property | Light reared  (24 cells from 7 mice) | Dark reared  (28 cells from 8 mice) | LMM (group effect) | |
| --- | --- | --- | --- | --- |
|  |  |  | F | P |
| Membrane potential (mV) | -78.3 ± 1.0 | -83.4 ± 1.0 | 13.03 | **0.004** |
| Input resistance (MΩ) | 121 ± 10.5 | 117 ± 9.88 | 0.002 | 0.96 |
| Membrane time-constant (ms) | 16.5 ± 1.29 | 16.5 ± 1.27 | 0.003 | 0.99 |
| Capacitance (pF) | 145 ± 6.81 | 139 ± 6.29 | 0.44 | 0.53 |
| Rheobase (pA) | 141 ± 10.7 | 119 ± 10.1 | 2.40 | 0.17 |
| Voltage threshold (mV) | -37.6 ± 0.796 | -38.7 ± 0.782 | 0.85 | 0.38 |
| AP amplitude (mV) | 79.3 ± 1.91 | 80.8 ± 1.90 | 0.29 | 0.60 |
| AP 20-80% rise-time (ms) | 0.14 ± 0.005 | 0.14 ± 0.005 | 0.26 | 0.63 |
| AP half-height width (ms) | 0.80 ± 0.03 | 0.88 ± 0.03 | 3.18 | 0.10 |
| AP maximum rise (mV.ms^-1^) | 408 ± 18.5 | 407 ± 18.2 | 0.003 | 0.96 |
| AP maximum decay (mV.ms^-1^) | 88.4 ± 4.26 | 80.6 ± 4.12 | 1.81 | 0.20 |
| IF Slope (AP.pA^-1^) | 0.049 ± 0.0035 | 0.046 ± 0.0034 | 0.23 | 0.64 |

Supplementary table 3: Summary of measured electrophysiological properties of recorded L2/3 neurons in V1B from dark reared mice.

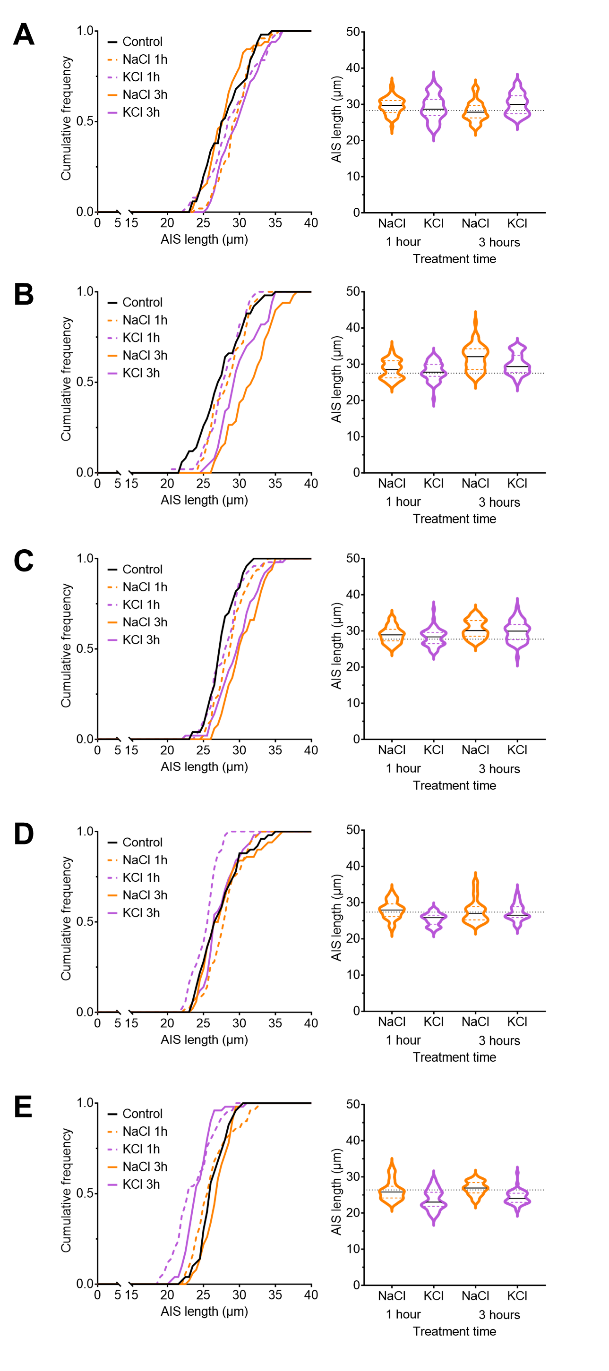
*Supplementary Figure 1: AIS lengths measured for each mouse included as part of this study.* Left, cumulative probability distributions for every mouse (**A-E**) measured for AIS length under control conditions (black lines), following 1-hour treatment with 15 mM NaCl (orange dashed lines and 15 mM KCl (purple dashed lines) or 3-hour treatment with 15 mM NaCl (orange solid lines and 15 mM KCl (purple solid lines). Right, violin plots of the same data, depicting the distribution of AIS lengths contributing to each mean shown in Figure 1C.

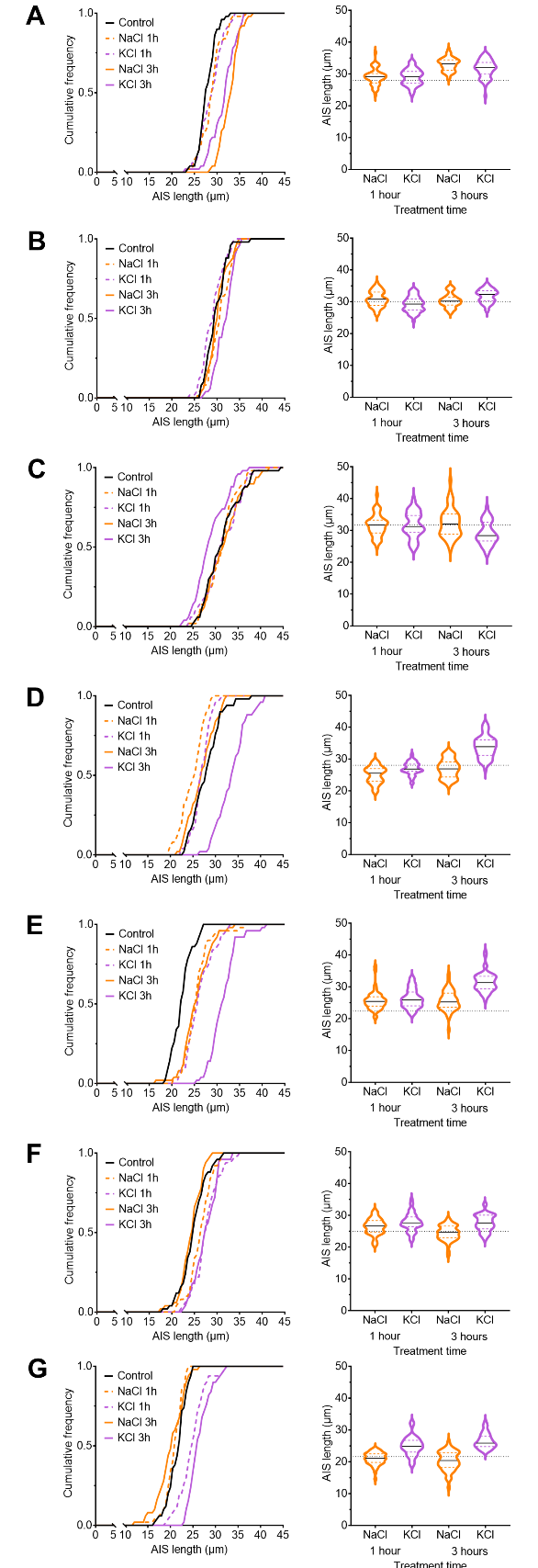

*Supplementary Figure 2: AIS lengths measured for each rat included as part of this study.* Left, cumulative probability distributions for every mouse (**A-E**) measured for AIS length under control conditions (black lines), following 1-hour treatment with 15 mM NaCl (orange dashed lines and 15 mM KCl (purple dashed lines) or 3-hour treatment with 15 mM NaCl (orange solid lines and 15 mM KCl (purple solid lines). Right, violin plots of the same data, depicting the distribution of AIS lengths contributing to each mean shown in Figure 1D.

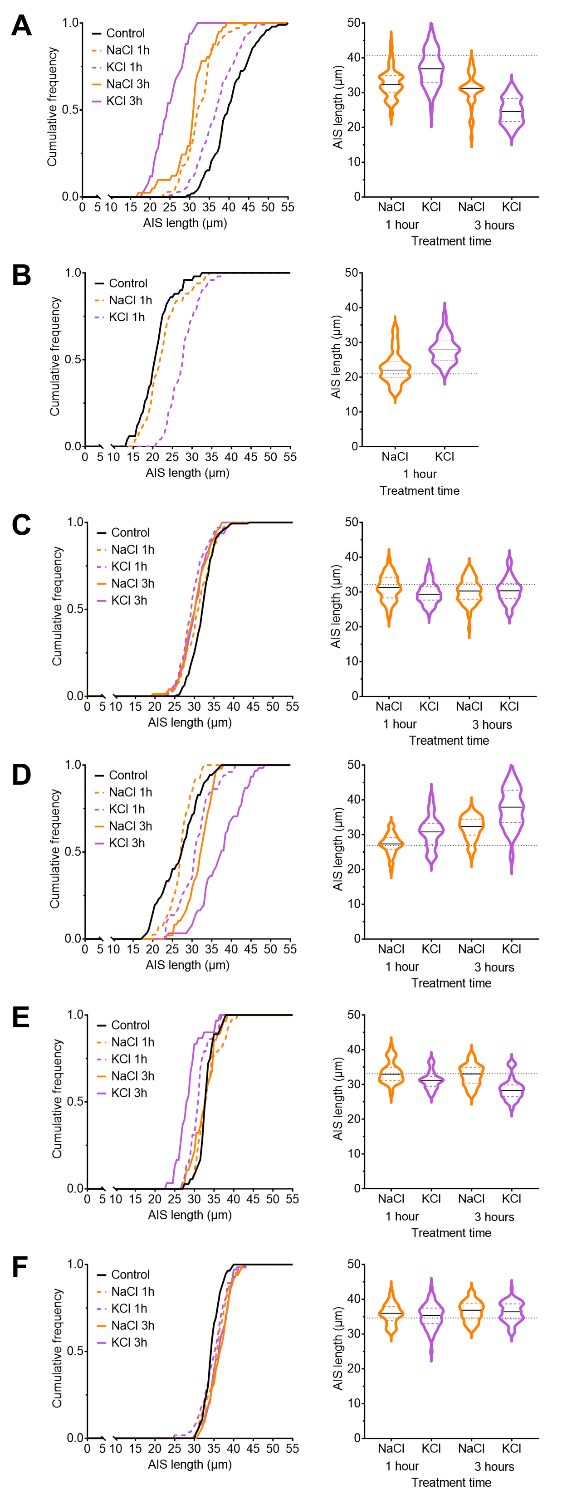

*Supplementary Figure 3: AIS lengths measured for each human cortical sample included as part of this study.* Left, cumulative probability distributions for every mouse (**A-E**) measured for AIS length under control conditions (black lines), following 1-hour treatment with 15 mM NaCl (orange dashed lines and 15 mM KCl (purple dashed lines) or 3-hour treatment with 15 mM NaCl (orange solid lines and 15 mM KCl (purple solid lines). Right, violin plots of the same data, depicting the distribution of AIS lengths contributing to each mean shown in Figure 1D.

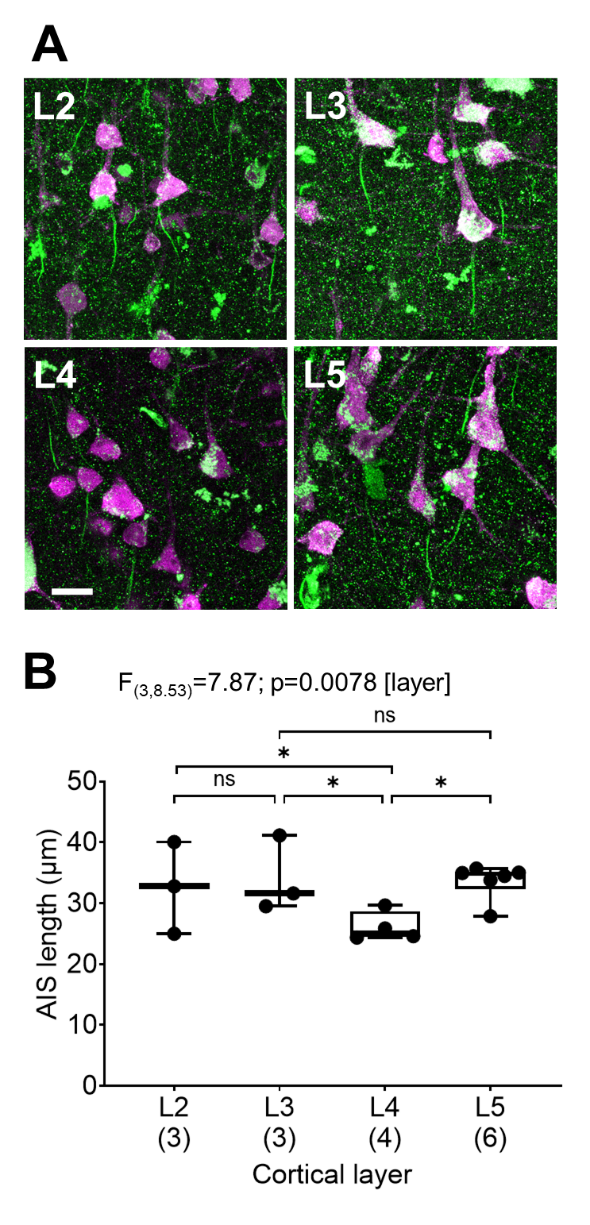

*Supplementary Figure 4: AIS lengths of neurons in different layers of human cortex.* A) example flattened confocal stacks of AnkyrinG (green) and NeuN (purple) labelling in L2, L3, L4, and L5 of human cortex. B) quantification of AIS length averaged per case for each layer. Average data from each patient are shown overlaying the mean ± SEM. Statistics shown from LMM.
